# Enhancer Identification using Transfer and Adversarial Deep Learning of DNA Sequences

**DOI:** 10.1101/264200

**Authors:** Dikla Cohn, Or Zuk, Tommy Kaplan

## Abstract

Enhancer sequences regulate the expression of genes from afar by providing a binding platform for transcription factors, often in a tissue-specific or context-specific manner. Despite their importance in health and disease, our understanding of these DNA sequences, and their regulatory grammar, is limited. This impairs our ability to identify new enhancers along the genome, or to understand the effect of enhancer mutations and their role in genetic diseases.

We trained deep Convolutional Neural Networks (CNN) to identify enhancer sequences in multiple species. We used multiple biological datasets, including simulated sequences, in vivo binding data of single transcription factors and genome-wide chromatin maps of active enhancers in 17 mammalian species. Our deep networks obtained high classification accuracy by combining two training strategies: First, training on enhancers vs. non-enhancer background sequences, we identified short (1-4bp) low-complexity motifs. Second, by replacing the negative training set by adversarial *k*-order random shuffles of enhancer sequences (thus maintaining base composition while shuttering longer motifs, including transcription factor binding sites), we identified a set of biologically meaningful motifs, unique to enhancers. In addition, classification performance improved when combining positive data from all species together, showing a shared mammalian regulatory architecture.

Our results demonstrate that design of adversarial training data, and transfer of learned parameters between networks trained on different species/datasets improve the overall performance and capture biologically meaningful information in the parameters of the learned network.

## 1 Introduction

An Accurate expression of genes is required for proper development and functioning of cells, and is largely achieved by regulating transcription. This regulation is mostly mediated by sequence-specific binding of proteins called transcription factors (TFs) to DNA regions called TF binding sites (TFBS). In this manner, a cell can regulate the transcription of many protein-coding genes (~20K in humans) using a relatively small number of TFs (8% of all proteins) (Narlikar et al., 2009).

Systematic computational and experimental approaches relying on RNA-sequencing and statistical properties of coding sequences enable automatic high-throughput annotation of genes in newly sequenced genomes, and protein-coding genes are widely investigated in model organisms and other species. In contrast, our understanding of regulatory elements controlling these genes is still limited. Current high-throughput experiments such as chromatin immunoprecipitation (ChIP-seq) test for in vivo binding across the genome, but are noisy, expensive, time-consuming and often available in few cell types and conditions. While less than 2% of the human genome encodes for proteins, it is estimated that ~5%-10% of genome is functional (Lindblad-Toh et al., 2011; Rands et al., 2014), and may contain non-coding regulatory elements. Experimental evidence (Wilson et al., 2008) shows that genome sequence largely determines the tissues and conditions under which such regulatory elements are active, thus motivating the development of computational approaches for identifying them using sequence data only *(de novo),* and for providing a better understanding of binding specificities and their interactions. To this end, we propose a computational model of deep **Convolutional Neural Network (CNN)** that learns to identify regulatory sequences in multiple species.

The most well-characterized types of regulatory elements are promoters and enhancers. While promoters are located near the transcription start site of genes, enhancers can regulate gene expression from afar, up to 1Mb away from their target gene (Stadhouders et al., 2012). Enhancers can also be located within introns or exons of other genes (Pennacchio et al., 2013); (Ahituv, 2016), do not necessarily affect the nearest gene, and may affect several genes (Mohrs et al., 2001); (Fukaya et al., 2016), in tissue or context-specific manner (Attanasio et al., 2014; Ong et al., 2011). In addition, most loci identified by Genome-Wide Association Studies (GWAS) lie far away from genes, and several in-depth experimental analyses have revealed that mutations in those regions disrupt enhancer activity (Colbran et al., 2017), supporting the view that enhancer mutations underlie many genetic diseases, and highlighting the bio-medical importance of enhancer sequences (Engel et al., 2016).

Chromatin marks are commonly used as indicators of an enhancer state, defined as the ability for an enhancer to increase expression of target genes, and are categorized into inactive, poised and active states. In particular, histone H3 lysine 27 is acetylated (H3K27ac) during activation. Thus, enhancers can be identified by mapping regions enriched for H3K27ac via ChIP-seq experiments (Creyghton et al., 2010).

Previous studies have shown that gene functionality can be modified by changing the enhancer sequence only. For example, the morphological disappearance of limbs in snakes is associated with sequence changes disrupting the function of a critical limb enhancer almost 1Mb away (Kvon et al., 2016). Another example is the impact of changes in the human ZRS sequence, which have been carried out in transgenic mice, where the single-nucleotide changes result in anterior-limb expression in abnormal positions during development, consistent with preaxial digit outgrowth (Masuya et al., 2007).

Methods for identifying Transcription Factor Binding Sites (TFBS) within regulatory DNA sequences often use Position Specific Scoring Matrices (PSSMs) (Stormo, 2000). These matrices score each offset in the input sequence and identify high-scoring occurrences of the motif. PSSMs assume independence of positions within the binding site, such that the total score (or energy) of the protein-DNA interaction is the sum over the matrix columns (Benos et al., 2002). Several studies challenged the validity of this assumption, and suggested to consider interdependent effects to better explain protein–DNA interactions (Man et al., 2001; Bulyk et al., 2002; Barash et al., 2003). TFBSs can also be learned *de novo* from biological sequences, using enrichment of motifs in a set of target biological sequences (e.g., promoter or enhancer regions) compared to a background set of sequences (e.g., non-enhancers genomic regions). Huggins et al. developed a method for discovery of discriminative motifs in two groups of sequences (DECOD) (Huggins et al., 2011). Ghandi et al. suggested using gapped *k-* mers (gkm-SVM) (Ghandi et al., 2014). In addition, a wide range of generative approaches to discovering motifs have been developed, from EM-based MEME algorithm (Bailey et al., 2006) to MEME-ChIP (Machanick et al., 2011), HOMER (Heinz et al., 2010) and others (Tran et al., 2014). Finally, Leslie et al. suggested a new sequence-similarity kernel, the spectrum kernel, for use with SVMs in a discriminative approach (Leslie et al., 2002).

We use a discriminative deep learning approach to identify enhancers vs. genomic background sequences, using Convolutional Neural Networks (CNNs). Deep CNNs have been successfully applied to computer vision, speech recognition, natural language processing, audio recognition, and lately also to bioinformatics, often achieving state-of-the-art performance in classification tasks. Several neural network approaches yielded promising results in enhancer prediction. Basset (Kelley et al., 2016) trained CNNs on accessible genomic sites mapped in different cell types by DNase-seq. DeepEnhancer (Min et al., 2017) trained the deep learning model on the FANTOM5 permissive enhancer dataset, and afterward on ENCODE cell type-specific enhancer datasets. DECRES (Li et al.,2016) used a feedforward neural network to distinguish between different kinds of regulatory elements, such as active enhancers and promoters. Alipanahi et al. showed that CNN models achieve excellent results on the TFBS prediction task and are scalable to a large number of genomic sequences (Alipanahi et al., 2015). Lanchantin et al. introduced new convolutional and recurrent neural network models that further improved TFBS predictive accuracy (Lanchantin et al., 2016).

Due to the natural role of TFBSs in biological regulatory sequences, most architectures use convolutional layers (Ching et al., 2017). While many models of TFBS prediction are similar to those used in computer vision and NLP, genomic applications of CNNs may benefit from exploiting properties specific to DNA sequence data. For example, motifs may appear on either forward or reverse strand. Special convolution models use this property to share parameters and achieve invariance to the reverse-complement operation, thus enabling motif discovery in both strands (Shrikumar et al., 2017). Multi-layered neural networks can use the intermediate layers to build up multiple layers of abstraction (Nielsen, 2015), and are thus attractive for modeling complex combinatorial gene regulation. For instance, the neurons in the first layer may learn to recognize simple patterns in the DNA sequence (similarly to PSSMs), the neurons in the second layer could then learn to recognize more complex patterns, such as spatial relations between pairs and triplets of motifs (as in gapped k-mer models), the third layer could recognize pairs of TFBSs that appear in proximity, and so on, thus capturing the complexity of biological regulatory sequences. More generally, these multiple layers of abstraction seem likely to give deep networks a compelling advantage in learning to solve complex pattern recognition problems (Benjio, 2009; Pascanu et al., 2014).

In this work, we investigate and improve several strategies for training CNNs to classify enhancer sequences. By training against adversarial negative data (with identical k-mer distribution, for *k ≤* 4), and by combining features learned on different adversarial datasets, our networks are able to learn realistic features (such as TFBSs or parts of such), leading to improved classification accuracy. In addition, while the number of available putative enhancers defined in a specific genome may limit our prediction accuracy, we show that combining samples from multiple species improve classification performance, and furthermore that learned CNNs can be transferred between different species, implying conservation of regulatory logic.

Finally, while traditional deep neural networks yield abstract models that are challenging to interpret, our model forces the learnable filters to converge into biologically meaningful and interpretable features, including the explicit modeling of TF binding sites and their wiring. Indeed, we show that our networks often learn filters resembling known motifs from the literature, demonstrating that examining the learned network can reveal biological insights into enhancer architecture. We show that our CNN-based training methods yield successful enhancer prediction for simulated datasets and real-life enhancers. Our methods are available in a python software package on GitHub.

## 2 Results

In section 2.1 we present the deep convolutional neural networks used throughout this paper. Section 2.2 analyzes simulated data of a single transcription factor binding. In Section 2.3, we analyze real-life transcription factor ChIP-seq data in five vertebrates. Finally, in Section 2.4 we analyze putative enhancer data (defined using distal H3K27ac peaks) in 17 mammals.

### 2.1 Deep Convolutional Neural Network for DNA sequences

We designed a deep neural network to identify regulatory regions (enhancers) from sequence. Briefly, each DNA sequence *s*i** of length *N* is represented in a one-hot encoding matrix *x_i_* of size 4 × *N.* where for each position *k = 1,…,N* and nucleotide *j* ∊ {*′A′, ′C′, ′G′, ′T′}* we have *x_i_* (*j, k*) = 1 if *s_i_* (*k*) = *j* and *x_i_* (*j, k*) = 0 otherwise. The matrix *x_i_* is used as input for the neural network. The true label of each sequence *y_i_* ∊ {0,1} is also represented in a one-hot encoding vector 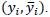

The general framework is illustrated in Figure 1. We train the deep network classifier using positive (enhancer) and negative (non-enhancer) sequences, possibly from various organisms, where training amounts to updating a parameters vector *W* which determines the mapping from input *x_i_* to output *y_i_*. We then test the performance of the trained network using held-out test data.

**Figure 1:**
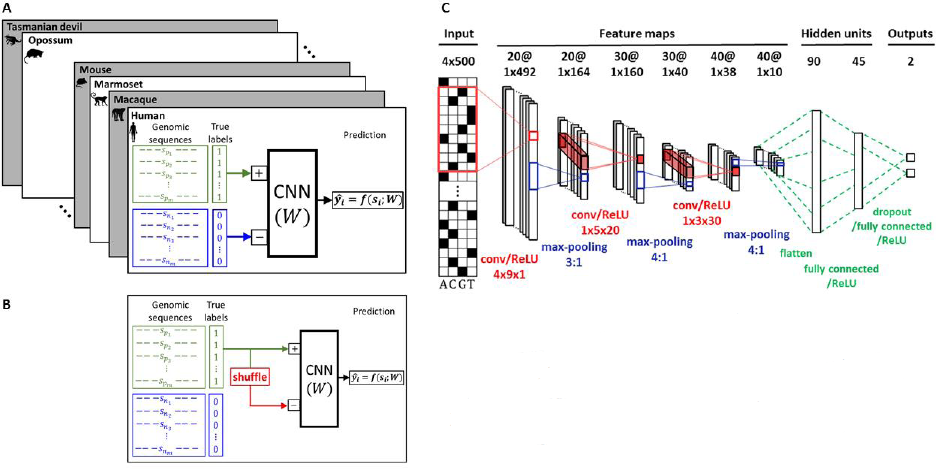
**A. Standard supervised training.** In the basic scenario, positive samples are regulatory regions (either bound by a single TF or enhancer sequences), and negative samples are non-enhancer background sequences. Networks are trained independently in multiple species. **B. Adversarial training.** The negative samples obtained by random shuffles of the positive sequences, thus maintaining their k^th^-other base composition. **C. Architecture of deep convolutional neural network (CNN) for enhancer prediction from DNA sequence**. Networks building blocks include convolutional, rectification, maxpooling, dropout and dense (fully connected) layers. The final output step uses a logistic function to predict that sequence label.

Positive sequences are defined as genomic regions showing in vivo binding by a single TF (for one set of sequences), or distal regions enriched for H3K27ac (for a second dataset). Negative (non-enhancer) sequences are composed of (1) distal genomic regions of length *N*, with no H3K27ac marks (Figure 1A); or (2) positive enhancer sequences that were randomly *k*-shuffled (Figure 1B), with a specific shuffling parameter *k* (See Section 2.3.4 and Methods, Section 3.1.3).

**The network architecture:** Briefly, our networks are composed of a series of convolutional layers. Each such layer has multiple filters, and is followed by ReLU and max-pooling steps. Finally, two fully connected layers are applied (via linear combinations), and the output layer predicts the (binary) label of the input DNA sequence (Figure 1C).

**Convolutional layers:** The first layers of the network are responsible for identifying sequence features along the input DNA sequence, such as TFBSs (or parts of such). To detect such features, during the training process we optimize a set of learnable filters (or kernels) that have a small receptive field, and extend through the full depth of the input volume. Each filter is convolved across the width and height of the input volume, computing the dot product between the entries of the filter and the input, and producing a 1-dimensional activation map of that filter. As a result, the network learns filters that are activated upon detecting a specific type of feature at some spatial position of the input.

**Reverse-complement:** A special convolutional function is applied in the first convolution layer of the CNN, to enable motif identification on both the forward and reverse strand by the same filter (see Methods).

**ReLU:** Following the convolution, a rectified linear unit (ReLU) is applied, zeroing-out negative inputs and keeping positive inputs intact.

**Max-pooling:** Smoothing is applied by taking the maximal value along consecutive non-overlapping windows, thus reducing the dimensionality of the feature map by a constant factor.

These stages are repeated several times, and are followed by two **fully connected layers,** where multiple linear combinations of the feature map are taken, and then a ReLU function is applied. To reduce *overfitting,* the second fully connected layer applies a **dropout** method, keeping each node of this layer with probability p = 0.85 at each training stage (Srivastava et al., 2014).

**Logistic layer**: Finally, a logistic function is applied in the final layer of the neural network, converting the two output scores into class probabilities at the output layer.

Convolutional neural networks generalize the classic sequence analysis methods: scanning a sequence using a PSSM can be viewed as a single (known) filter, followed by max-pooling of the entire sequence (yielding the single maximum score), which is then compared to a threshold value to classify the sequence.

**Model training:** We randomly split the input data into training, validation, and held-out test data (80%:10%:10%, for both positive and negative sets). We used the **cross-entropy** (logistic) loss – ubiquitous in modern deep neural networks – to compare the predicted scores vs. the true labels, for optimizing the model parameters. We used TensorFlow, with an Adaptive Moment Estimation (Adam) optimizer (Kingma et al., 2015) as a stochastic Gradient-based optimization algorithm, with mini-batches of 50 sequences for 20 epochs or until convergence (Methods).

**Model evaluation:** We evaluated our models’ performance on three tasks: (1) classifying simulated sequences (with or without planted motifs); (2) identifying DNA sequences bound by a single TF; and (3) identifying enhancer sequences enriched for *bona fide* binding sites of multiple TFs (2 and 3 in real-life biological context). We tested the performance of the learned networks in terms of the classification success rate, using the area under a ROC curve (AUC measure), and in terms of the biological insights distilled from the learned network parameters. To gain such insights, we examined the filters of the first convolutional layer (those scanning the DNA sequence itself), and compared them to known binding motifs from JASPAR (Sandelin et al., 2004) using the HOMER suite (Heinz et al., 2010) (see Methods).

### 2.2 Simulated Data

We started with a simple synthetic dataset, where we sampled 10K positive and 10K negative 500bp long DNA sequences from a background uniform distribution, with probability ¼ for each nucleotide. For positive samples, we planted a single “binding site” according to the One Occurrence Per Sequence (OOPS) model. Binding sites were sampled from the known Position Weight Matrix (PWM) of CEBPa, and planted at position *J,* according to a Normal distribution, *J~N*(250,40^2^) (see Methods). We then examined the performance of our network (with the architecture described in Figure 1C) on the simulated data. This network was chosen for the main task of enhancer classification (see Section 2.4).

We compared the results of our neural network to those of an oracle PSSM model, which uses the true PWM of CEBPa and positional distribution of sites. This oracle model has the lowest possible error (also known as Bayes' classification error), and serves as a gold standard for the classification task. For a competitive straw-man model, we used the PSSM model with the top *de novo* motif identified using HOMER tool (Heinz et al., 2010), with or without a positional bias.

As shown in Figure 2, our CNN (blue; AUC=0.81) outperforms the straw-man models (AUC=0.74, 0.69 with or without positional bias, respectively). The performance of our CNN is reasonably close to that of the oracle model (AUC=0.9), and better than an oracle model with no positional bias (AUC=0.79). These results suggest that the CNN managed to learn both sequence preferences (PWM) and the distribution of planted binding site locations. The performance gap with respect to the oracle can be partly explained by the limited sample size. As sample size increases to 30K samples, our model is within 0.035 from the oracle AUC, indicating that with a large enough sample size, our network may achieve near-oracle performances.

**Figure 2:**
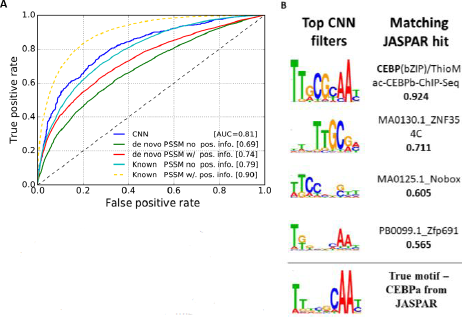
**A. ROC curve of different models, trained and tested on simulated CEBPa ChIP-seq data.** CNN performance (blue) is compared to PSSM models with known PWM (cyan/yellow) or following *de novo* motif discovery (green/red); with or without explicit modeling of binding site positional information (using a Normal distribution). Yellow line marks oracle performance.**B. Learned filters match planted CEBPa motif**. We compared all 20 1 ^st^-layer filters learned by the CNN to known JASPAR motifs (using HOMER similarity score), and ranked them by their similarity to the closest JASPAR motif. The top four CNN filters match the known CEBPa motif (including half-motif and variants).

### 2.3 Single TF: CEBPa and HNF4A binding in the liver

We next tried to identify regulatory regions enriched with a specific type of TF binding site using real biological sequences. We defined positive samples using peaks called from ChIP-seq for two liver transcription factors: CEBPa and HNF4A (Schmidt et al., 2010), (Supp. Info., Table 1). We next describe the analysis of CEBPa binding in five vertebrates: Human, Mouse, Dog, Opossum, and Chicken. HNF4A yielded qualitatively similar results (Supp. Info., Section 2.1). To avoid bias, we extracted an equal number of 12K positive samples for each species. The network described above (Figure 1C) performed well when running on this classification the TF data (average AUC of 0.93), but **it failed to learn meaningful biological motifs** in the first layer of filters. We therefore tried different network architectures and chose a CNN with two convolutional layers (average AUC of 0.92, see Figure 3), where the first layer has seven filters of size 4×9×1 each, and the second layer has 50 filters of size 1×2×7 each.

**Figure 3:**
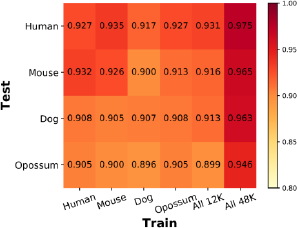
Multi-species learning improves CEBPa ChIP-seq classification. Each column represents a trained network (12K positive and negative samples), and each row shows test accuracy (ROC I AUC) in different species. Similar accuracy is reported regardless of training species. Rightmost column shows improved accuracy when combining training samples from all species (48K positive/negative samples).

#### 2.3.1 TF ChIP-seq peaks vs. genomic background sequences

We begin by using biological non-enhancer sequences as negative data. For each species, we sampled a random set of genomic regions (500 bp long), excluding regions with N’s, regions less than 15Kb from genes (Zerbino et al., 2018), and H3K27ac bound regions (Villar et al., 2015). The latter data was not available for chicken which we excluded from this analysis. We trained our CNNs on each species separately, with 12K positive and 12K negative samples. The results, shown in Figure 3 on diagonal, show successful enhancer prediction in all four species with similar AUC, all in the range 0.9-0.92.

#### 2.3.2 Transfer Learning Between Different Species

We next asked to what extent can samples from one species improve our network performance in other species. Such transferability can have practical consequences as it allows us to predict enhancers in species with limited experimental data (or none whatsoever). More importantly, it can reveal to what extent is regulatory logic shared between species.

We trained our network using samples from all species together (48K positive and 48K negative). For a fairer comparison with the single-species results, we also considered a dataset with 3K positive and 3K negative samples from each species.

As shown in Figure 3, networks trained on different species are transferable – and allow training on one species and testing in another, thus showing the conservation of a regulatory code (at least for CEBPa). Moreover, training on data from multiple species together dramatically improved the test accuracy vs. single-species training (AUC increased from 0.9-0.92 to 0.95-0.97; Figure 3, rightmost column), thus reducing the error rate by over 50%.

#### 2.3.3 Biological interpretation of the learned filters

As shown in Figure 3, our CNN performs very well in the classification task of TF binding sites vs. non-enhancer negative data. A closer examination (Figure 4) shows that it failed to learn the “correct” motif, and instead learned low-complexity DNA motifs that discriminate between TF peaks and non-enhancer background. Indeed, regulatory sequences were shown to contain short repetitive motifs – mostly GC rich – that promote their activity (Yáñez-Cuna et al., 2014, Colbran et al., 2017).

**Figure 4:**
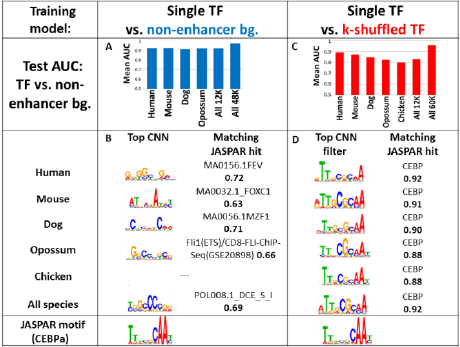
CNN results on TF ChIP-seq peaks vs. background (non-enhancer) sequences, after training vs. naive (left) or adversarial (right) samples. **A**. Average AUC for held-out test data (ChIP-seq vs. non-enhancer sequences) for networks trained on different species. **B**. Top motifs learned by the CNN model trained against non-enhancer background samples. All CNN filters were compared to all JASPAR motifs (using the HOMER score), and the top hits are shown. **C.** Same as (A), for CNNs trained against adversarial (*k*-shuffled) samples. **D**. Same as (B), for CNN trained against adversarial (*k*-shuffled) samples. Here, all filters identified the true CEBPa motif (bottom) as the highest scoring PWM. **Despite the marginally better classification of CNNs trained against non-enhancer background samples (A-B), these CNNs failed to learn biologically significant motifs, suggesting that a combination of adversarial and transfer learning should be applied.** Only low complexity motifs were learned by filters of the first training model (A), and they did not resemble the CEBPa motif.

To improve our model's accuracy and to learn more complex, biologically meaningful filters, we employed an adversarial approach. Specifically, we shuffled the positive sequences to generate negative samples that are identical in their *k*^th^-order base composition to positive samples, thus forcing our network to focus on discriminative motifs longer than k, instead of short low-complexity motifs (Figure 1B).

#### 2.3.4 Training against *k*-shuffled adversarial sequences

**The negative adversarial examples** define a different and possibly harder classification problem, compared to our original problem of classifying regulatory regions vs. non-enhancers. As we increase *k*, the negative samples become more similar to the positive ones, making the classification task harder.

**Choosing k**: To determine *k*, we trained our CNN to discriminate between genomic *(non-enhancer)* sequences (here serving as positive examples) and their *k*-shuffled form (negative examples) for different values of *k.* As shown in Figure 5, we observed a dramatic deterioration in AUC up to *k* = 4. After which, the positive sequences are almost indistinguishable from their *k*-shuffles (AUC≤0.55). After choosing *k* (using *non-enhancer* sequences), we use the same *k* for comparing positive sequences and their *k*^th^-order shuffles, thus focusing on features that are unique to the ChIP-seq peaks and are not due to general genomic composition. At *k* = 4, ChIP-seq peaks are still easily separated than their *k*-shuffles (AUC=0.9, compared to 0.55 of the non-enhancers vs. their *k*-shuffles).

**Figure 5:**
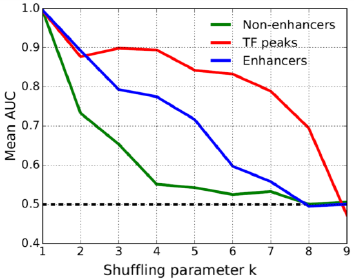
Average classification accuracy (shown as mean AUC) at various k values. Shown are (1) comparison of negative non-enhancer sequences vs. their *k*-shuffles (green); CEBPa ChIP-seq peaks vs. their *k*-shuffles (red); and active enhancers (marked by H3K27ac) vs. their *k*-shuffles (blue). All models were trained and tested on samples from all species together. Y-axis marks the test ROC AUC, average over all species. Identification of shuffled non-enhancer sequences becomes hard at *k*=4 (AUC=0.55). Conversely, ChIP-seq sequences are still separable from their shuffles (using sequence features longer than *k*, AUC=0.9). Intriguingly, enhancer sequences seem to contain a combination of short and long features, leading to a gradual decrease in accuracy as *k* is increased.

**Adversarial samples improve interpretation:** When training on TF ChIP-seq peaks vs. their k-shuffles *(k* = 4), some of the convolutional filters in the first layer of the network converged into a motif similar to the known PWMs. Each filter was compared to all JASPAR PWMs and the top filter per species is reported, together with the known CEBPa motif from JASPAR (Figure 4). In all species, CEBPa was identified as the top motif, with high HOMER similarity score (>0.88) (Heinz et al., 2010), compared to (<0.72) similarity for the network previously trained (Section 2.3.1) vs. non-enhancer sequences. To conclude, by replacing the negative set with random *k*-order shuffles of regulatory sequences, we explicitly learn biologically meaningful motifs.

### 2.4 Classification of Enhancer Sequences

We next turned to our main problem of identifying enhancers. For this purpose, we trained our CNNs on active enhancer sequences defined as active enhancer marks (H3K27ac) in liver across 17 mammalian species from Villar et al., 2015 (see Methods and Supp. Table 2). We extracted 14K positive and negative samples for each species.

#### 2.4.1 Enhancers vs. biological negative data

We first trained our CNNs to discriminate between enhancers and biological non-enhancer sequences, using 14K positive and 14K negative examples in each species.

We tried different hyper-parameters and CNN architectures when working with the enhancer data against non-enhancers, adjusting the number of convolutional layers, number of filters in each convolutional layer and size of each filter, number of fully connected layers and size of each layer, etc. This makes sense since enhancers may involve multiple types of TF binding sites compared to the single-TF case. For example, when running the previous network used for the single TF on the enhancer data, it yielded bad results (AUC=0.58). We finally chose the network detailed in Figure 1C for the enhancer classification, containing three convolutional layers, first one with 20 filters of size 4×9×1 each, second with 30 filters of size 1 × 5×20 each, and third with 40 filters of size 1×3×30 each. This network performed well on the enhancer vs. non-enhancer classification task, as shown in Figure 7.

**Figure 7:**
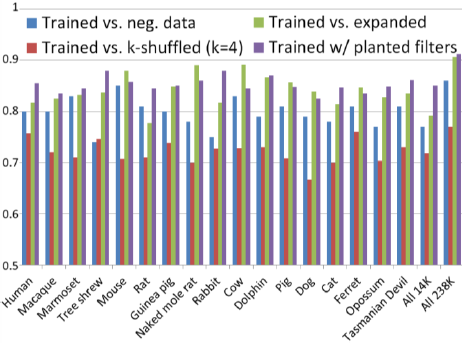
Transfer and Adversarial learning contribute to better enhancer classification. Shown are classification results of enhancer vs. background (nonenhancer) sequences, after training at different scenarios, including: (1) enhancers vs. non-enhancer sequences (blue), (2) enhancers vs. *k*-shuffled enhancer sequences (red), (3) networks trained on enhancers vs. expanded negative data (non-enhancers and *k*-shuffled data), and (4) networks trained on enhancers vs *k*-shuffled data, with filters then “planted” into new larger models. All models were tested on enhancer (H3K27ac) vs. background (non-enhancer) sequences. We also tried to use a larger CNN (with the same number of parameters as the CNN with planted filters), where all parameters are randomly initialized This network did not improve the result of the original CNN trained on enhancer vs. non-enhancer sequences (the larger randomly initialized network gives an AUC of 0.79, while the original smaller randomly initialized network gives an AUC of 0.80).

As in the case of a single TF, training of enhancer sequences vs. *bona fide* non-enhancer sequences yields low-complexity DNA motifs, rather than longer functional binding sites. Our trained model captures the background base composition (e.g., dinucleotide distribution) discriminating enhancers from the biological non-enhancer background.

Supp. Figure 3 shows classification accuracy for cross-species training. In similar to the single TF case, the learned network models are transferable between species. Test accuracy on each species of the best network – the one trained on 238K positive enhancer sequences and 238K negative non-enhancer biological sequences from all species together, is lower (average AUC=0.86 vs. 0.95) compared to the case of a single TF, in agreement with our expectation of complex combinatorial regulatory logic involving multiple TFs controlling enhancers, making their identification harder compared to identifying singe-TF binding sites. We next employed adversarial training, proven successful for the single TF problem, to improve our performance for the problem of identifying enhancers.

#### 2.4.2 Histone Modifications vs. *k*-shuffled negative data

We trained our network to discriminate between the positive enhancer sequences (H3K27ac peaks) and their negative random shuffles with varying *k*-order Markov model. We also examined whether here too, the network will learn biologically meaningful filters, especially in the first convolutional layer, corresponding to liver TF binding motifs. Since we do not know in advance all the possible binding motifs defining liver enhancers, we searched the learned filters against known motifs, and recorded their similarity scores (Heinz et al., 2010) and information content. For this step, all motifs in JASPAR are used for comparison.

Overall, the similarity and information content were lower, compared to the case of single TF, but higher compared to the non-adversarial networks.

#### 2.4.3 Transfer Learning Between Different Species

Like the single TF case, we examined the transferability of samples between species. To this end, we trained our model on samples from all species together (238K positive and 238K negative), and on a subset of samples from all species, containing a portion of the samples of each species (total of 14K positive and 14K negative).

Similarly to the single TF case, networks trained on different species are exchangeable, and training on data from multiple species together yielded better results compared to training on each species individually (see Supp. Figure 3). Moreover, transferability here reflected phylogeny, as Opossum and Tasmanian devil, the only two marsupial mammals in our dataset, had significantly lower classification test accuracy, compared to the fifteen eutherian mammals.

#### 2.4.4 Improving Classification with Adversarial Training

Using the insights gained during training with different background sets, we devised two strategies in which the adversarial approach is used to improve our classification performance at the original task of discriminating enhancer vs. non-enhancer sequences, described next.

##### 2.4.4.1 Expanded negative data

In the first approach, we expanded the negative training data, to consist of two types of negative sets: The first part contains *k*-shuffled enhancer sequences, with different values of *k*, and the second part contains biological non-enhancer sequences. We tried different partitions of these negative sets, and then chose the best partition. For the first part, we enumerated over all 2^9^ possible subsets of *k =* {1, …,9}. For the second part, we created 3 different sets by duplicating the biological negative data *n* = 1,2 or 3 times. We trained vs. all 2^9^ x 3 combinations using 10 runs, discarded all combinations yielding AUC<0.6 in their best run, and used 50 runs for the remaining combinations. The negative set achieving the best results is shown in Figure 6A, with biological negative data duplicated twice *(n =* 2), thus comprising 40% of the negative samples, and *k*-shuffling taken for *k =* 2,3,4, each comprising 20% of negative samples. The positive samples have been duplicated 5 times, in order to have a balanced training set with equal number of positive and negative examples.

**Figure 6:**
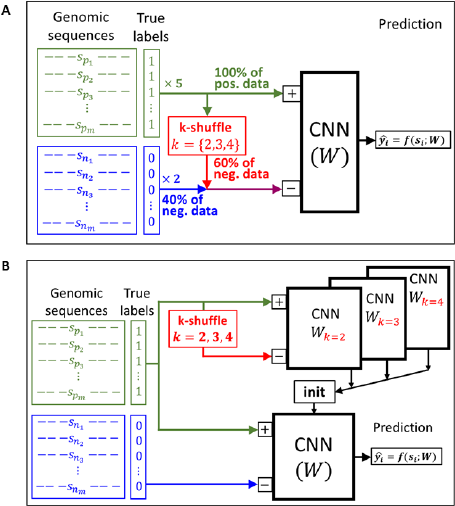
Improved network. **A.** Diagram of CNN trained on expanded negative data, 40% containing biological negative data and 60% containing *k*-shuffled enhancer sequences with *k*= {2,3,4}. **B.** Diagram of CNN with ‘planted’ filters. 60% of the convolutional layers’ parameters were initialized according to previous networks’ parameters – networks that were trained against *k*-shuffle of the positive enhancer sequences, with *k*= {2,3,4}. The rest of the convolutional layers’ parameters (40%) and the fully connected parameters were randomly initialized.

##### 2.4.4.2 CNN with ‘planted’ filters

In this approach, we used the filters learned by the adversarial training, rather than the adversarial examples themselves. We initialized some of the CNN's parameters by 'planting' filters learned by the adversarial trained networks, trained on enhancers vs. *k*-shuffles for *k =* 2,3,4, as opposed to random parameter initialization. Specifically, the filter parameters in all 3 convolutional layers are now initialized according to filter parameters learned earlier by one of the three adversarial trained networks (for *k =* {2,3,4}) and the fourth part of parameters were randomly initialized. All parameters in fully connected layers were also randomly initialized.

This training strategy is illustrated in Figure 6B. As shown in Figure 7, these two methods indeed improved the CNN performance in the enhancer vs. non-enhancer classification task, with average AUC increasing from 0.80 to 0.86. While negative data expansion is a simpler and more general approach, smart initialization by filter planting achieved slightly better performance (AUC=0.86 vs. AUC=0.845).

## 3 Methods

### 3.1 Deep Convolutional Neural Network Model

Our deep convolutional neural network represents a mapping *ŷ_i_* = Pr(*y_i_* = 1) = *f*(*s_i_ ; W*) from the sequence *s_i_* (or its one-hot encoding representation *x_i_*) into class probabilities. We denote by *W* the learnable parameters of the network, including all weights and biases. We used the TensorFlow Python library (ver. 1.1.0) to design, train and test the CNN model, and scikit-learn for generating ROC curves and AUC measure.

**Reverse complement convolutions:** To account for reverse complement occurrences of each filter, we added a special convolutional function in the first layer of the CNN (Shrikumar et al., 2017). At every position in the input sequence, we applied each filter in both orientations, and chose the maximum one as output for the next layer. This was done by reversing the first two dimensions (rows, columns) of each filter, equivalent to rotating the matrix that represents the filter by 180 degrees. Consecutive layers are therefore invariant to the strand in which a neuron was activated.

**Logistic transformation:** The two output neurons of the network *z_0_, z_1_*(Figure 1) are converted into positive class probability by applying the logistic function 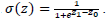

#### 3.1.1 Training the CNN Model

**Cross-entropy loss function:** Consider *N* labeled training samples 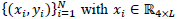 the sequence one-hot representation and *y_i_* ∈ {0,1} the label for sample *i.* The cross-entropy loss of a network parameterized by *W* for a sample *(x_i_,y_i_)* is denoted by *l(W; X,y_i_),* and is given by:

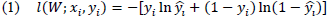

Where 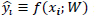 is the predicted value of the model. We optimize *W* by minimizing the loss function over all samples: L(W;X,y) =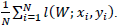

#### 3.1.2 Parameter optimization

The network parameters *W* (weights and biases) are trained using the Adaptive Moment Estimation (Adam), a stochastic Gradient-based optimizer (Kingma et al., 2015). Briefly, this algorithm uses a momentum-based gradient approach to iteratively update *W.*

The update rule is based on a subset of training examples – *mini-batch.* At each iteration *t* we choose a mini-batch *B_t_ ⊂ {1,…, N}* of size *b,* and define the loss of this iteration as:

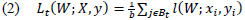

Adam uses the stochastic gradient over the mini-batch *∇L_t_(W_t_;X,y)* to estimate the first two moments of the gradient (parameter-wise) at the current parameterization *W_t_*. Adam then uses these estimates to normalize the gradient by dividing its norm by its mini-batch estimated standard deviation, giving the update rule at iteration *t*: (assuming tolerance parameter 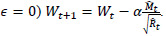 See Supp. Info for a full description. We used Adam’s default hyper-parameters: the learning rate *α=* 0.001, the 1^st^ moment decay rate *β_1_ = 0.9,* the 2^nd^ moment decay rate *ß_2_ = 0.999,* and the tolerance parameter *∊ =* 10^−8^. We used mini-batches of size *b =* 50. In each training epoch we go through all the training samples, using 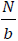 non-overlapping mini-batches, and shuffle them at the end of each epoch. In each run, we performed up to 20 learning epochs 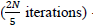-we allowed early stopping by calculating the train and validation accuracy every 2 iterations (100 samples), and stopped if the mini-batch loss was extremely low *(L_t_(W;X,y) < 0.0002).* We performed 50 runs of training (with random initializations of the network parameters *W),* and saved the model with maximal validation accuracy over all runs.

**Comparison of motif matrices:** We examined the filters of the first convolutional layer that scan the DNA sequence itself, and compared it to known motifs from the JASPAR database (Sandelin et al., 2004) using HOMER v4.6 (Heinz et al., 2010). Briefly, HOMER similarity reflects the Pearson correlation between a query PWM and the best JASPAR match (after trying all possible alignments, including reverse-complements).

#### 3.1.3 Adversarial Negative Data

**Generating k-shuffled background sequences**: Background sequences were generated using a *k*-order shuffle of (positive) enhancer sequences using varying *k* values (Kandel et al., 1996; Jiang et al., 2008). Specifically, we iteratively identified a *k* bp-long motif (k-mer) that occurs several times within each sequence, and randomly shuffled the sub-sequences between them (Figure 8). This procedure generates a random shuffle of the original sequence, while maintaining the exact number of occurrences for each *k*-mer within each sequence. Algorithmically, this can be done by constructing a de Bruijn multigraph in which the vertices **{v_i_}** represent the k-mers appearing in the original sequence s, and edges correspond to (*k*+ 1)-mers in the sequence, thus connecting two (k-mer) nodes, with an overlap of *k-1* bases. Random k-order shuffles of the original sequences can be then obtained by finding an Eulerian path in this graph. We used the uShuffle tool implementing this algorithm (Jiang et al., 2008).

**Figure 8:**
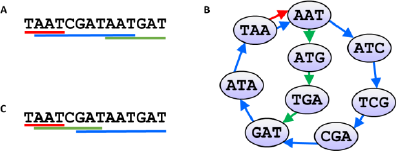
*k*-order shuffling represents alternative Eulerian paths in the *k*-order de Bruijn multigraph. **A.** An original genomic sequence for demonstration. **B.** The corresponding de Bruijn multigraph of order 3**. C.** A new possible shuffled sequence, with the same triplet counts as in the original sequence.

#### 3.1.4 Simulated Data

We simulated data using a simple One Occurrence Per Sequence (OOPS) model: each positive sample contains a single occurrence of the binding site, sampled (base by base) from multinomial distributions according to a given Position Weight Matrix (PWM). The short sub-sequence is planted in a random background sequence, with bases sampled i.i.d, from a uniform distribution. The location of the planted sub-sequences is normally distributed around the center of the long sequence *(µ = 250, σ^2^ = 40^2^),* resembling real data from noisy ChIP-seq experiments (Ma et al., 2014). Negative samples were sampled similarly, with no planted binding sites added. We downloaded JASPAR PWMs for two known motifs of liver-specific transcription factors: CEBPa (MA0102.2) and HNF4A (MA0114.1) (Sandelin et al., 2004). We then added pseudo-counts of 0.01 to avoid matrix entries having a value of 0 and re-normalized the PWMs. For each of the two PWMs, we created a synthetic data with 10K positive samples and 10K negative samples.

### 3.2 Single TF Data in Livers of Five Vertebrates

We next turned to test our method in real biological settings, where the goal is to identify regulatory regions enriched for a specific type of transcription factor binding site. We analyzed the genome-wide binding of two liver-specific transcription factors: (1) CEBPa in five vertebrates: human, mouse, dog, opossum and chicken, and (2) HNF4A in human, mouse, and dog (Schmidt et al., 2010). For both TFs, we used peaks called from Chromatin-immunoprecipitation sequencing (ChIP-seq) data to define bound loci in each species (Supp. Tablel). We defined a 500bp window centered at each peak, and used the bedtools package (Quinlan et al., 2010) to extract the corresponding DNA sequences as our positive TF samples.

Here we describe the CEBPa analysis. A similar analysis for HNF4A is shown in the Supp. Info. In order to have the same amount of samples from each species, we extracted the positive samples corresponding to 12K peaks for each species.

We trained our CNN on each species separately (12K positive and 12K negative samples), on all species excluding Chicken (48K positive and 48K negative samples), and on a random subset of samples from all species, containing one fourth of the samples of each species (total of 12K positive and 12K negative samples).

### 3.3 Enhancer Sequences: H3K27ac Peaks in Livers of 17 Mammals

We next turned to predict enhancer sequences that contain *bona fide* binding sites for multiple TFs, in real biological context. We used data from Villar et al., who tracked the evolution of promoters and active enhancers across the livers of 20 mammalian species (Villar et al., 2015). Of those, we excluded three species due to lack of genome sequence or poor annotations (Supp. Table2). To avoid bias due to differences in the number of called peaks, we extracted 14K peaks as positive samples for each species.

We then trained our CNN on each species separately (14K positive and 14K negative samples), on all species together (17 × 14 = 238K positive and 238K negative samples), and on a random subset of samples from all species (total of 14K positive and 14K negative).

### 3.4 PSSM model

The PSSM straw-man model attempts to separate the positive and negative samples by giving each sample a score, based on the best match to an input PWM.

Briefly, for a sequence *s* of length *N* and a PWM *P*^(*M*)^ of length *L*, we assume the OOPS model, where a specific single location is chosen at random for the motif, according to a prior *P*^(*L*)^ We also have a uniform background model *P^(B)^.* In this model, we compute the Maximum Log-Likelihood Ratio (MLLR) statistic over all possible positions (see Supp. Info.):

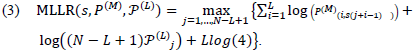

We then compare it to a fixed threshold. For a model with no positional bias, the score above simplifies to the standard PSSM scoring. In addition to the known PWMs, we also used the highest scoring *de novo* motif, as identified using HOMER's findMotifs function (Heinz et al., 2010).

## 4 Discussion

By incorporating both biological non-enhancer sequences and *k*-order adversarial shuffles of enhancer sequences, we captured and used both low complexity features (representing background base composition), as well as longer features (such as DNA binding motifs for regulatory proteins). As we showed, combining these two strategies improved our ability to discriminate between enhancers and non-enhancers from DNA sequence data only, thus advancing our understanding of the underlying regulatory grammar.

Naïve application of deep-learning methods to DNA sequence in biological settings may achieve satisfactory classification performance. However, it is hard to biologically interpret the resulting parameters of trained network. As a consequence, these models do not advance our understanding of the underlying biological grammar of DNA motifs and the molecular mechanisms involved.

In contrast, training deep neural networks on carefully designed adversarial data, forces the network to learn a meaningful data representation, thus harnessing the computational power of deep learning into biological insights.

Specifically, we developed two such strategies: first, training against negative heterogonous dataset comprising a mixture of negative samples and adversarial *k*-shuffled positive samples. Second, by a sequential approach where network filtered learned using one dataset were planted into a network trained on another dataset.

We have shown the utility of transferring learned filters between different species. Future work may improve transferability between various species by utilizes their evolutionary relationships. In addition, the transfer-learning methodology has far reaching implication beyond the scope of this problem. For example, we can use enhancers from one cell/tissue-type to improve enhancer prediction on another cell/tissue-type.

Our work opens the door for new possibilities in interpreting deep networks applied in genomic context. We have shown that filters learned at the first convolutional layer have a clear biological interpretation as short motifs and known TF binding sites. By carefully examining additional layers we propose to interpret more complex, higher-order regulatory interactions between various mechanisms of gene regulation.

## Acknowledgements

This research was funded by the Israeli Center of Excellence (I-CORE) for Gene Regulation in Complex Human Disease (no. 41/11), by a Marie Curie Career Integration Grant (PCIG13-GA-2013-618327), and by an Israel Science Foundation grant (no. 913/15) to T.K.

## References

Ahituv, N. (2016). Exonic enhancers: proceed with caution in exome and genome sequencing studies. Genome Medicine, 8(14).

Alipanahi, B., Delong, A., Weirauch, M. & Frey, B. (2015). Predicting the sequence specificities of DNA- and RNA-binding proteins by deep learning. Nat Biotechnol., 33(8), 831–838.

Attanasio, C., Nord, A., Zhu, Y. et al. (2014). Tissue-specific SMARCA4 binding at active and repressed regulatory elements during embryogenesis. Genome Res., 24(6), 920–929.

Bailey, T., Williams, N., Misleh, C. & Li, W. (2006). MEME: discovering and analyzing DNA and protein sequence motifs. Nucleic Acids Res., 34(Web Server issue), W369–W373.

Barash, Y., Elidan, G., Friedman, N. & Kaplan, T. (2003). Modeling Dependencies in Protein-DNA Binding Sites. Proceedings of the 7th annual international conference on Computational molecular biology, 28–37.

Benjio, Y. (2009). Learning Deep Architectures for AI. Foundations and Trends in Machine Learning, 2(1), 1–127.

Benos, P., Bulyk, M. & Stormo, G. (2002). Additivity in protein-DNA interactions: how good an approximation is it? Nucleic Acids Res., 30(20), 4442–4451.

Bulyk, M., Johnson, P. & Church, G. (2002). Nucleotides of transcription factor binding sites exert inter-dependent effects on the binding affinities of transcription factors. Nucleic Acids Res., 30(5), 1255–1261.

Ching, T., Himmelstein, D., Beaulieu-Jones, B. et al. (2017). Opportunities and obstacles for deep learning in biology and medicine. bioRxiv, doi: 10.1101/142760.

Colbran, L., Chen, L. & Capra, J. (2017). Short DNA sequence patterns accurately identify broadly active human enhancers. BMC Genomics, 18(536).

Creyghton, M., Cheng, A., Welstead, G. et al. (2010). Histone H3K27ac separates active from poised enhancers and predicts developmental state. Proc. Natl. Acad. Sci. (PNAS). 107(50), 21931–21936.

Engel, K., Mackiewicz, M., Hardigan, A., Myers, R. et al. (2016). Decoding transcriptional enhancers: Evolving from annotation to functional interpretation. Semin. Cell Dev. Biol., 57, 40–50.

Fukaya, T., Lim, B. & Levine, M. (2016). Enhancer Control of Transcriptional Bursting. Cell, 166(2), 358–368.

Ghandi, M., Lee, D., Mohammad-Noori, M. & Beer, M. (2014). Enhanced Regulatory Sequence Prediction Using Gapped k-mer Features. PLoS Comput. Biol., 10(7), 1–15.

Heinz, S., Benner, C., Spann, N. et al. (2010). Simple Combinations of Lineage-Determining Transcription Factors Prime cis-Regulatory Elements Required for Macrophage and B Cell Identities. Mol. Cell, 38(4), 576–589.

Huggins, P., Zhong, S., Shiff, I. et al. (2011). DECOD: Fast and Accurate Discriminative DNA Motif Finding. Bioinformatics, 27(17), 2361–2367.

Jiang, M., Anderson, J., Gillespie, J. & Mayne, M. (2008). uShuffle: a useful tool for shuffling biological sequences while preserving the k-let counts. BMC Bioinformatics, 9(192).

Kandel, D., Matias, Y., Unger, R. & Winkler, P. (1996). Shuffling biological sequences. Discrete Applied Mathematics, 71(1-3), 171–185.

Kelley, D., Snoek, J. & Rinn, J. (2016). Basset: learning the regulatory code of the accessible genome with deep convolutional neural networks. Genome Res., 26(7), 990–999.

Kingma, D.P. & Ba, J.L. (2015). Adam: A Method for Stochastic Optimization. International Conference for Learning Representations.

Kvon, E.Z., Kamneva, O.K., Melo, U.S. et al. (2016). Progressive loss of function in a limb enhancer during snake evolution. Cell, 167(3), 633–642.

Lanchantin, J., Singh, R., Wang, B. & Qi, Y. (2016). Deep Motif Dashboard: Visualizing and Understanding Genomic Sequences Using Deep Neural Networks. arXiv, 1608.03644.

Leslie, C., Eskin, E. & Noble, W. (2002). The spectrum kernel: a string kernel for SVM protein classification. Pac. Symp. Biocomput., 564–575.

Li, Y., Shi, W. & Wasserman, W.W. (2016). Genome-Wide Prediction of cis-Regulatory Regions Using Supervised Deep Learning Methods. bioRxiv, doi: 10.1101/041616.

Lindblad-Toh, K., Garber, M., Zuk, O. et al. (2011). A high-resolution map of human evolutionary constraint using 29 mammals. Nature, 478(7370), 476–482.

Ma, W., Noble, W. & Bailey, T. (2014). Motif-based analysis of large nucleotide data sets using MEME-ChIP. Nat. Protoc., 9(6), 1428–1450.

Machanick, P. & Bailey, T. (2011). MEME-ChIP: motif analysis of large DNA datasets. Bioinformatics, 27(12), 1696–1697.

Man, T. & Stormo, G. (2001). Non-independence of Mnt repressor-operator interaction determined by a new quantitative multiple fluorescence relative affinity (QuMFRA) assay. Nucleic Acids Res., 29(12), 2471–2478.

Masuya, H., Sezutsu, H., Sakuraba, Y. et al. (2007). A series of ENU-induced single-base substitutions in a long-range cis-element altering Sonic hedgehog expression in the developing mouse limb bud. Genomics, 89(2), 207–214.

Min, X., Zeng, W., Chen, S. et al. (2017). Predicting enhancers with deep convolutional neural networks. BMC Bioinformatics, 18(Suppl 13):478, 35–46.

Mohrs, M., Blankespoor, C., Wang, Z. et al. (2001). Deletion of a coordinate regulator of type 2 cytokine expression in mice. Nature Immunology, 2, 842–847.

Narlikar, L. & Ovcharenko, I. (2009). Identifying regulatory elements in eukaryotic. Brief. Funct. Genomic. Proteomic., 8(4), 215–230.

Nielsen, M. (2015). Neural Networks and Deep Learning. Determination Press. Retrieved from http://neuralnetworksanddeeplearning.com.

Ong, C. & Corces, V. (2011). Enhancer function: new insights into the regulation of tissue-specific gene expression. Nat. Rev. Genet., 12(4), 283–293.

Pascanu, R., Montufar, G. & Bengio, Y. (2014). On the number of response regions of deep feedforward networks with piecewise linear activations. arXiv, 1312.6098.

Pennacchio, L., Bickmore, W., Dean, A. et al. (2013). Enhancers: five essential questions. Nat. Rev. Genet., 14(4), 288–295.

Quinlan, A. & Hall, I. (2010). BEDTools: a flexible suite of utilities for comparing genomic features. Bioinformatics, 26(6), 841–842.

Rands, C., Meader, S., Ponting, C. & Lunter, G. (2014). 8.2% of the Human genome is constrained: variation in rates of turnover across functional element classes in the human lineage. PLoS Genet., 10(7), e1004525.

Sandelin, A., Alkema, W., Engström, P. et al. (2004). JASPAR: an open-access database for eukaryotic transcription factor binding profiles. Nucleic Acids Res., 32(Database issue), D911–D94.

Schmidt, D., Wilson, M., Ballester, B. et al. (2010). Five-vertebrate ChIP-seq reveals the evolutionary dy-namics of transcription factor binding. Science, 328(5981), 1036–1040.

Shrikumar, A., Greenside, P. & Kundaje, A. (2017). Reverse-complement parameter sharing improves deep learning models for genomics. bioRxiv, doi: 10.1101/103663.

Srivastava, N., Hinton, G., Krizhevsky, A. et al. (2014). Dropout: A Simple Way to Prevent Neural Networks from Overfitting. Journal of Machine Learning Research 15(1), 1929–1958.

Stadhouders, R., van den Heuvel, A., Kolovos, P. et al. (2012). Transcription regulation by distal enhancers. Transcription, 3(4), 181–186.

Stormo, G. (2000). DNA binding sites: representation and discovery. Bioinformatics, 16(1), 16–23.

Tran, N. & Huang, C. (2014). A survey of motif finding Web tools for detecting binding site motifs in ChIP-Seq data. Biology Direct, 9(4).

Villar, D., Berthelot, C., Aldridge, S. et al. (2015). Enhancer evolution across 20 mammalian species. Cell, 160(3), 554–566.

Wilson, M., Barbosa-Morais, N., Schmidt, D. et al. (2008). Species-specific transcription in mice carrying human chromosome 21. Science, 322(5900), 434–438.

Yáñez-Cuna, J., Arnold, C., Stampfel, G. et al. (2014). Dissection of thousands of cell type-specific enhancers identifies dinucleotide repeat motifs as general enhancer features. Genome Res., 24(7), 1147–1156.

Zerbino, D.R., Achuthan, P., Akanni, W. et al. (2018). Ensembl 2018. Nucleic Acids Res., 46(Database issue), D754–D761.

